# The biosynthetic gene cluster landscape of the oral microbiome across health and dental caries

**DOI:** 10.1101/2025.09.04.674288

**Authors:** McKenna Loop Yao, Peijun Lin, Kailey Hua, Wenjun Zhang

## Abstract

Specialized metabolites encoded by biosynthetic gene clusters (BGCs) in the oral microbiome remain largely unexplored in the context of oral health and disease. Previous genome-centric surveys have cataloged hundreds of uncharacterized BGCs in the oral cavity associated with health and disease, but these studies relied on reference genomes and did not capture strain-level variation or the native distribution of BGCs. Here, we assembled three independently sourced metagenomic datasets from healthy and dental caries samples, extracted BGCs, and quantified their abundance alongside expression in a metatranscriptomic dataset. We first identified that aryl polyene, ribosomally synthesized and post-translationally modified peptide (RiPP), and nonribosomal peptide (NRP) encoding BGCs were the most abundant BGC classes across all three metagenomic datasets. We then grouped these BGCs into homology-based families and found that homologous clusters were usually consistently associated with either health or dental caries, suggesting conserved community-level roles for BGCs. An elastic-net regression model further selected 45 BGCs out of >5000 that could distinguish healthy and dental caries samples in the metatranscriptomic dataset, which demonstrated that BGCs could be predictive markers of disease. This analysis emphasizes the importance of high-quality metagenomic and metatranscriptomic datasets to resolve BGC expression patterns and to guide discovery of metabolites relevant to oral health and disease.

## Introduction

Dental caries is one of the most prevalent diseases worldwide (Shoaee et al., 2024). Unlike traditional one-pathogen models of infection, dental caries arises from local microbial dysbiosis, which is characterized by a shift from a diverse, health-associated microbial community to a less diverse community dominated by acidogenic and aciduric species that promote demineralization of the tooth surface (Bostanghadiri et al., 2024; X. Li et al., 2022; Spatafora et al., 2024; Zhu et al., 2023). Early studies on dental caries often relied on 16S rRNA gene sequencing and emphasized taxonomic shifts in microbial communities associated with disease (Willis & Gabaldón, 2020). However, disease-associated taxa are often inconsistent across individuals and populations, making solely taxonomic analysis insufficient for characterizing the onset of disease (Moussa et al., 2022). Recent omics-based research has suggested that microbial functional activity might be a more reliable marker of pathogenesis, with specific biochemical markers, such as carbohydrate metabolism and acid production, found to shape community structure and clinical outcomes (Duran-Pinedo & Frias-Lopez, 2015; Wang et al., 2019). Thus, a critical next step in understanding and preventing caries involves the comprehensive characterization of the specific biochemical drivers of oral pathogenesis (Anwer et al., 2025; Qadri et al., 2024).

Specialized metabolites (SMs) are emerging as crucial biochemical contributors to human health and disease, having been found to be involved in mediating interspecies competition, modulating virulence, enhancing metal acquisition, and enhancing biofilm formation (Barber & Zhang, 2021; Donia et al., 2014). SMs are a diverse group of small molecules, encompassing non-ribosomal peptides (NRPs), polyketides (PKs), ribosomally synthesized and post-translationally modified peptides (RiPPs), terpenoids, saccharides, alkaloids, and various hybrids or derivatives of primary metabolites (Newman & Cragg, 2020). Any particular microbial SM is typically produced via a dedicated biosynthetic gene cluster (BGC), which includes co-localized biosynthetic genes on a microbial genome (Walsh & Fischbach, 2010). These BGCs encode characteristic biosynthetic enzymes that can be readily predicted from genomic information via BGC predictors, such as antiSMASH or DeepBGC (Blin et al., 2025; Liu et al., 2022).

Previous genome-centric surveys have identified several hundred uncharacterized BGCs in the oral cavity associated with health or disease by mining public genomes and mapping them to metagenomes (Aleti et al., 2019; Koohi-Moghadam et al., 2024). However, these studies did not capture the native prevalence of BGC classes within the oral microbiome or investigate strain-level differences in BGC landscape. To overcome these limitations, we directly extracted BGCs from three independently sourced, metagenomic datasets associated with dental caries and oral health. We then analyzed these BGCs’ prevalence in health and disease metagenomic samples and, in parallel, correlated their abundance within a metatranscriptomic dataset to assess *in situ* activity. We finally predicted the BGCs’ species origins and elucidated trends in BGC homology and correlation with health or disease. This approach allowed us to capture global trends in BGC presence and expression across independent datasets. Notably, we show that quantitative BGC presence and expression can diverge from their species’ presence in dental-caries status, and BGCs can serve as predictors of oral health status. Together, these insights underscore the need for integrated, high-resolution whole-genome metagenomic and metatranscriptomic studies to obtain complete functional profiles of the oral microbiome and prioritize BGC-driven discovery.

## Results

Integrating our metagenomic-assembly workflow, we first obtained three previously published metagenomic datasets (Datasets 1-3) associated with dental caries and oral health. Each sample was assembled with metaSPADES (Nurk et al., 2017), and its BGCs were predicted with antiSMASH 7.0 (Blin et al., 2023). A custom Python script then removed duplicate BGCs (>90% similarity) within a given dataset (see Methods), which yielded a non-redundancy BGC list for each dataset. These non-duplicate BGCs were aligned back to the raw (unassembled) samples with BWA-MEM2 (Vasimuddin et al., 2019), and per-sample counts were generated to compare BGC prevalence between health and caries conditions. Despite differences in cohort origin, extraction protocols, and sequencing platforms, the metagenomic-assembly and subsequent BGC identification workflow revealed consistent trends in BGC representation across all three datasets, underscoring its robustness for cross-study comparisons.

### BGC landscape of Dataset 1

Dataset 1 comprised whole-genome metagenomic samples of saliva from 44 preschool children (ages 45 to 73 months), which included 25 with severe early childhood caries (ECC) and 19 age-matched, caries-free controls (Wang et al., 2019). Metagenomic DNA libraries were prepared using the Illumina TruSeq DNA sample prep v2 guide and sequenced on a HiSeq2000 instrument. Their analysis found 20 species enriched in the severe ECC group, which included multiple *Prevotella* spp., *Streptococcus mutans,* and lesser studied strains *Prevotella amnii*, *Shuttleworthia satelles*, *Olsenella uli*, and *Anaeroglobus geminatus*. They also found that healthy samples exhibited increased abundance of *Neisseria lactamica* or *Streptococcus australis.* Their metabolic functional analysis also revealed that the dental caries ECC group exhibited a significant enrichment in functions related to sugar metabolism (Wang et al., 2019).

Application of the metagenomic-assembly workflow to Dataset 1 yielded 3,636 biosynthetic gene clusters. Removal of redundant BGCs with our dereplicate script resulted in 1,378 BGCs, of which the core biosynthetic genes were then taxonomically annotated via BLASTp (Camacho et al., 2009) and NCBI’s ClusteredNR database (default database as of 2025) and default BLASTp parameters (see Methods). Four BGC families made up nearly 70% of the total extracted BGCs, which included RiPP-like, aryl polyene, NRP and NRP-like, and RiPP Recognition Element (RRE)-containing BGCs (**Fig. 1A**). BGCs were then mapped to metagenomic samples using bwa-mem2, and alignments with a BWA-MEM2 score greater than 80 were counted. Counts of BGCs in caries and healthy samples were compared using the Wilcoxon rank-sum test, and BGCs with *p* < 0.05 were considered significantly associated with either dental caries or health. This resulted in 125 BCGs significantly associated with health and 97 with dental caries (**SI Datasheet 1, Tab ‘Dataset 1’**). Among these significant BGCs, we found that terpene BGCs were more enriched in healthy samples and NRP BGCs were more enriched in caries samples (**Fig. 1A**). Next, we identified the 20 BGCs with the largest positive Cohen’s *d* values (standardized mean differences between health and caries), which represent the BGCs most enriched in caries samples (**SI Datasheet 2**). Interestingly, nearly all of these 20 BGCs originated from *Prevotella* species (e.g. *Prevotella histicola, Prevotella sp.* HMSC073D09, *Prevotella nigrescens*) and encoded aryl polyene, RiPP-like, β-lactone, and resorcinol biosynthetic pathways. Conversely, the 20 BGCs with the most negative Cohen’s *d* values were dominated by *Neisseria* species and included primarily aryl polyene, RiPP-like, and terpene classes (**SI Datasheet 2**). These results suggest that disease-associated BGCs are commonly derived from disease-associated species, and health-associated BGCs predominantly derive from health-associated species.

**Figure 1:**
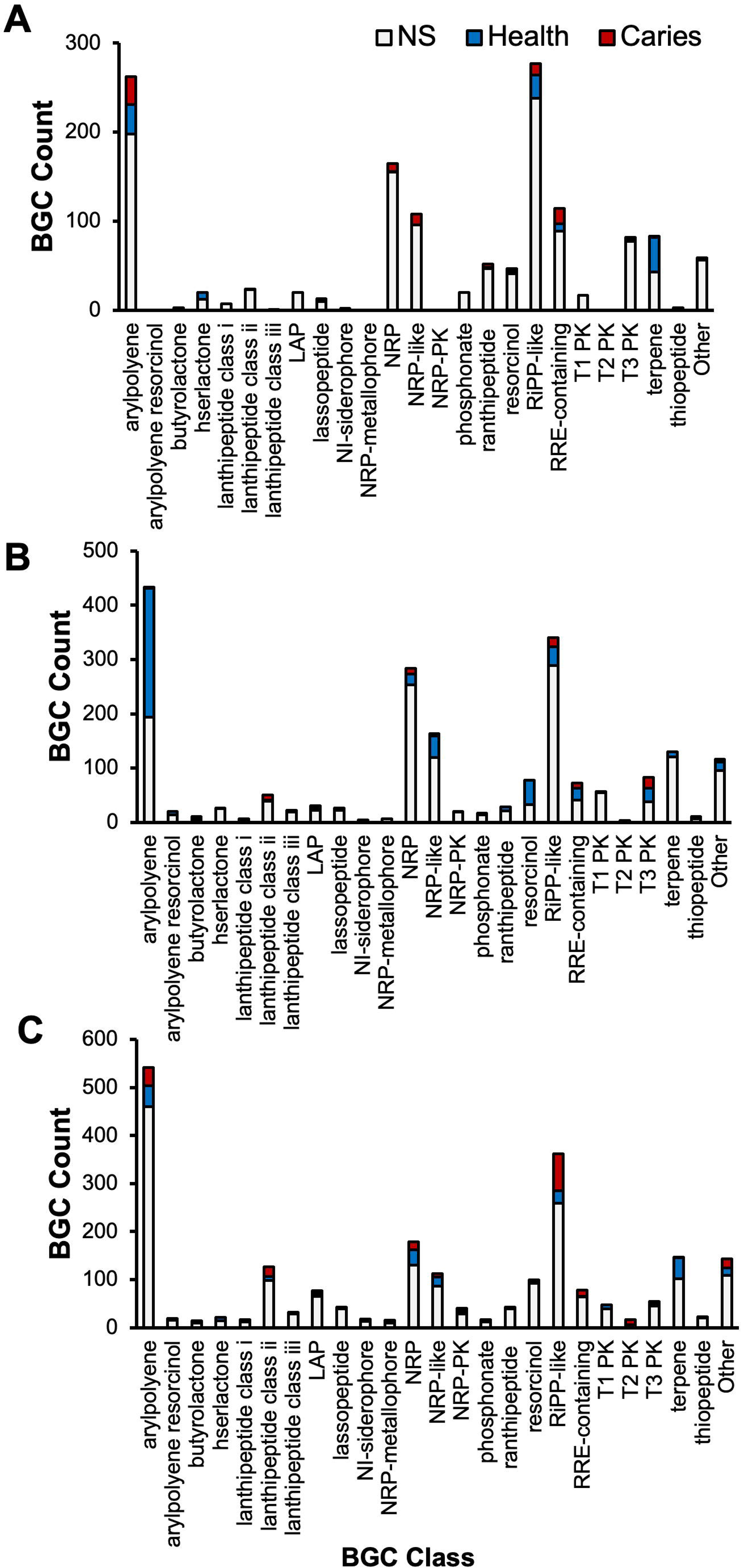
Biosynthetic gene cluster (BGC) class distributions in Datasets 1-3. Bar plots show the distribution of BGC classes in Datasets 1-3 (A-C, respectively). Significant caries­ associated BGCs are colored red, health-associated BGCs in blue, and non-significant (NS) BGCs in white.

We next performed a focused examination comparing the species-level taxa reported in Dataset 1’s original taxonomic analysis with the BGC correlations identified in the present study. We identified 21 RiPP-like BGCs from various strains of *Neisseria lactamica*, a species correlated with health in the original taxonomic analysis; however, only 11 of this species’ BGCs were significantly associated with health (**SI Datasheet 1, Tab ‘Dataset 1’**). In another health-correlated species, *Streptococcus australis*, we identified five RiPP-like BGCs, only two of which were associated with health. Additionally, Dataset 1’s original taxonomic analysis also reported that *Neisseria mucosa* was present in both healthy and caries samples and therefore part of the natural oral microbiota, but we found several terpene BGCs from *Neisseria mucosa* that were associated with health in our workflow (**SI Datasheet 1, Tab ‘Dataset 1’**). Dataset 1’s original taxonomic analysis also reported *Prevotella buccae, Prevotella amnii, Prevotella sp.* oral taxon 317 and *Prevotella oris* were enriched in caries samples. We found no BGCs in these reported species; instead, we identified several BGCs (arlypolyene, RiPP-like, thiopeptide, β-lactone), including many caries associated ones, in other *Prevotella* species (e.g. *Prevotella histicola*, *Prevotella sp.* HMSC073D09, *Prevotella nigrescens*, and *Prevotella pallens*) (**SI Datasheet 1, Tab ‘Dataset 1’**). These results demonstrate that health- and disease-associations at the BGC level can diverge from those observed at the species level.

Lastly, we clustered the 1,378 BGCs into gene-cluster families (GCFs) with BiG-SCAPE (Navarro-Muñoz et al., 2020), which compares BGC pairs by Pfam domain composition, domain order, and backbone gene sequence identity to group loci that likely encode homologous BGCs (**Fig. 2., S1**). We visualized the GCFs’ association with oral health and disease on Cytoscape (Shannon et al., 2003) and color-coded the BGCs (nodes) on a gradient from red (caries) to blue (health) corresponding to the magnitude of their Cohen’s *d* values. From this analysis, we observed many GCFs that were confined predominantly within one species, suggesting the species-specific metabolite family and biological functions. In contrast, several GCFs contained BGCs from several species, supporting possible broad ecological roles. Also, the BGCs within a given GCF usually exhibited the same association with either health or caries, suggesting that these homologous BGCs could support similar ecological roles. For example, a RiPP-like GCF that contained BGCs from various strains of *Neisseria lactamica* and a terpene GCF with BGCs from *Neisseria* and *Kingella* species were both associated with health (**Fig. 2**). Conversely, a multi-*Prevotella* spp. and *Segatella salivae* aryl polyene GCF and a mixed *Prevotella/Segatella* RRE-containing GCF were consistently linked to caries (**Fig. 2**). Interestingly, while the *Prevotella/Segatella* GCF also included homologous RRE-containing gene clusters from various *Hoylesella* species, none of these *Hoylesella* BGCs were associated with disease (**Fig. 2**). This suggests that homologous BGCs from different hosts may have diverged to yield chemically distinct metabolites with unique bioactivities, or they may still produce the same compound whose effect shifts with the surrounding microbial and host context. Therefore, meaningful interpretation of BGC-phenotype relationships may need to consider both the possible structural diversity of metabolites within GCFs and the ecological context in which their products operate.

**Figure 2:**
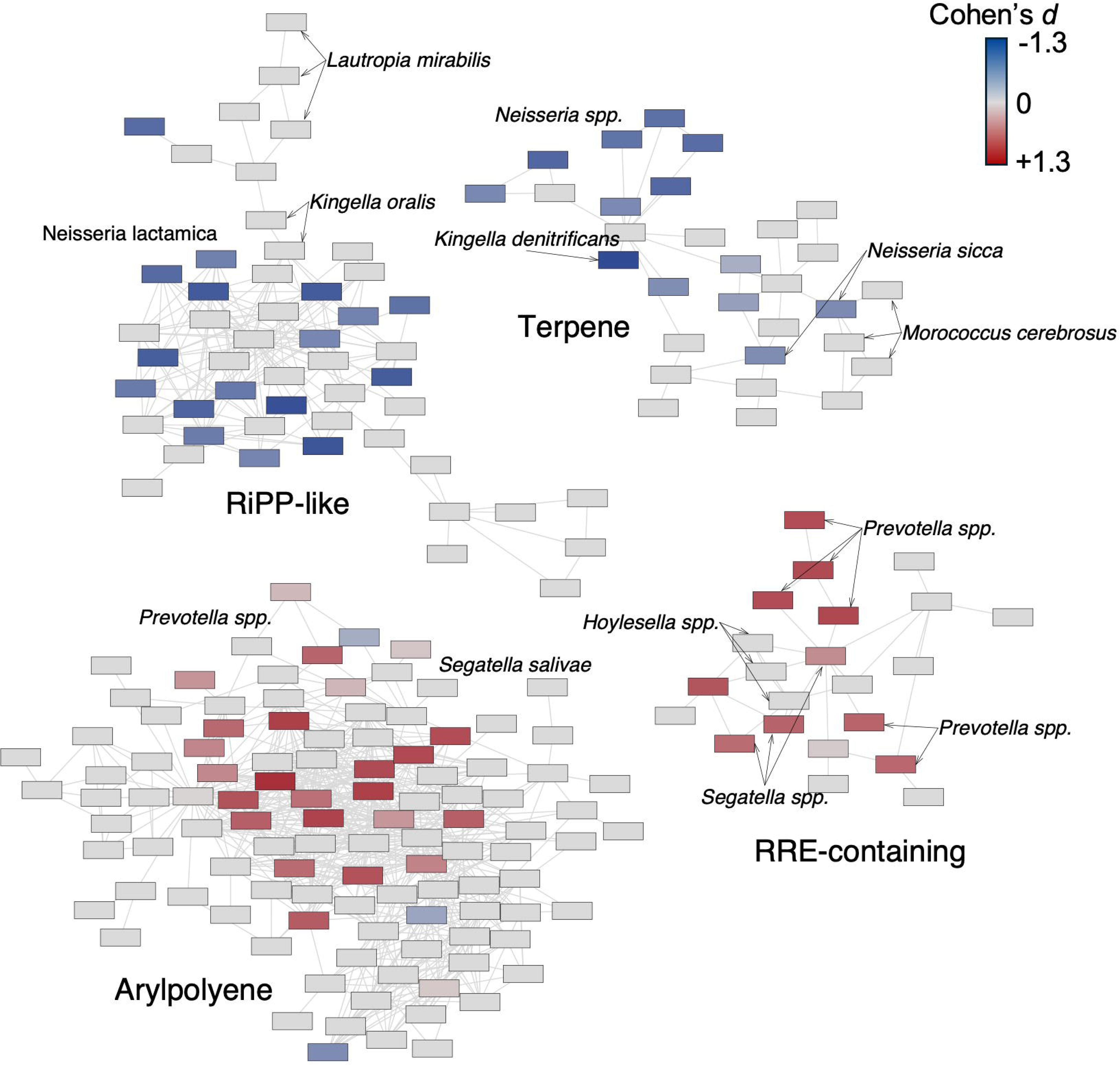
Selected BiG-SCAPE gene cluster families (GCFs) from Dataset 1. Selected gene cluster families from Dataset 1 are shown; see Fig S1 for complete GCF map for Dataset 1. GCFs were grouped using BiG-SCAPE (v2.0) with Pfam-A as the domain database, automatic alignment mode, inclusion of singletons, and a GCF similarity cutoff of 0.5. The resulting similarity network was visualized in Cytoscape using an organic layout, with edges displayed only for distances between 0.3 and 1.0 and filtered to include edge similarity scores (DSS) between 0.5 and 1.0. Nodes represent single biosynthetic gene clusters (BGCs), and edges represent pairwise similarities above the specific thresholds. The GCFs are labeled as their dominant BGC class, and the color gradient indicates the magnitude of each BGC’s Cohen’s *d* value, with blue denoting stronger associations with health and red denoting stronger associations with caries.

### BGC landscape of Dataset 2

Dataset 2 included 31 healthy and 57 dental caries metagenomic samples of dental plaque samples from kids aged 5 to 11 years old (Espinoza et al., 2018). Plaque was collected by thoroughly swabbing gingival margins and buccal surfaces. Metagenomic libraries were sequenced using the Illumina NextSeq 500 High-Output kit for 300 cycles. Taxonomically, *Streptococcus*, *Catonella morbi*, and *Granulicatella elegans* were enriched in caries-positive subjects, while *Tannerella forsythia*, *Gracilibacteria*, *Capnocytophaga gingivalis*, *Bacteroides sp. oral taxon 274*, and *Campylobacter gracilis* were more abundant in the healthy cohort. Functionally, 37 KEGG metabolic modules were positively associated with caries-positive microbiomes, including numerous phosphotransferase sugar uptake systems and an increased abundance of two-component histidine kinase-response regulator systems. Their analysis concluded that caries onset coincides with the perturbation of an entire ecosystem and is characterized by enhanced sugar catabolism rather than a single bacterium (Espinoza et al., 2018).

Application of the metagenomic-assembly workflow on Dataset 2 generated 5,800 candidate biosynthetic gene clusters (BGCs) which reduced to 2,036 non-redundant BGCs post dereplication. Consistent with Dataset 1, aryl polyenes, RiPP-like/RRE-containing loci, and NRP/NRP-like BGCs made up the majority of BGCs extracted from the assembled metagenomes (**Fig. 1B**). Different from Dataset 1, Dataset 2 displayed a marked bias toward health-associated BGCs, with 489 significantly associated with health and 91 associated with disease (Wilcoxon rank sum test *p* < 0.5) (**SI Datasheet 1, Tab ‘Dataset 2’**). More than half of all aryl polyene BGCs were significantly enriched in health samples, whereas only two aryl polyene BGCs exhibited a caries association (**Fig. 1B**). The resorcinol pathways that frequently co-occurred with the aryl polyene BGCs showed the same health bias (45 health; 0 caries) (**Fig. 1B**). The 20 BGCs with the largest effect size in the direction of caries based on highest Cohen’s *d* values included a lanthipeptide-class-ii/lassopeptide loci from *Rothia aeria*, an RRE-containing cluster from *Actinomyces*, an NRPS-like pathway in *Rothia aeria*, an NRPS pathway from *Streptococcus mutans*, and several RiPP-like clusters from *Streptococcus oralis* and *Streptococcus pneumoniae* (**SI Datasheet 2**). In contrast, the 20 most health-associated BGCs were dominated by aryl polyene pathways in multiple *Capnocytophaga* species, including oral taxon 863, *leadbetteri*, *granulosa*, and oral taxon 335. In agreement with Dataset 1, Dataset 2 showed that the BGCs most strongly linked to health or caries resided in species with matching associations. However, the BGCs that held the highest magnitude of Cohen’s *d* values between the two datasets differed, indicating that the strength of BGC correlations can vary with extraction method, sampling site, and cohort composition.

We next compared Dataset 2’s original taxonomic analysis with the present study’s BGC health and disease associations. Dataset 2’s original analysis identified several *Streptococci* species (including *S. sanguinis, S. mutans,* and *S. mitis*) enriched in caries samples, and we similarly found a RiPP-like BGC and a type III PK BGC from *S. mitis* and several lanthipeptides from *S. sanguinis* that all showed a weak but significant caries association (**SI Datasheet 1, Tab ‘Dataset 2’**). However, *S. mitis* also harbored a RiPP-like BGC that we found to be weakly associated with health (Cohen’s *d* = 0.16), underscoring the value of BGC-level analysis for functional insight, rather than sole reliance on taxonomic analysis. Current limitations in genome availability and assembly methods hinder accurate strain-level classification in taxonomic analyses, often obscuring unique pathogenic traits. Different strains can possess markedly divergent pangenomes, which have been shown to vary in their ability to contribute to health or disease. Our findings indicate that BGC-based analysis can resolve certain strain-level differences without the need for extensive taxonomic characterization or large-scale genome sequencing.

We next integrated BiG-SCAPE networking to organize the 2,036 BGCs into discrete GCFs and colored the BGC nodes based on their Cohen’s *d* values (**Fig. 3, S2**). Two large *Capnocytophaga* spp. Aryl polyene GCFs, along with a large mixed-species aryl polyene GCF containing Aggregatibacter spp., Cardiobacterium sp., Campylobacter sp., and Neisseria spp., were uniformly health-associated, emphasizing the broad prevalence of aryl polyene BGCs among health-associated taxa (**Fig. S2**). Additionally, *Leptotrichia* spp., *Fusobacterium vincentii*, and *Fusobacterium polymorphum* each harbored a health-associated GCF with NRP and NRP-like BGCs, suggesting that health-associated species also encode more specialized metabolic pathways (**Fig 3**). On the disease side, two GCFs comprised both *Rothia aeria* and *Rothia dentocariosa*: one NRP GCF and one lanthipeptide class-II GCF. Markedly, both GCFs displayed contrasting species disease associations. Specifically, in the NRP GCF, BGCs from *Rothia dentocariosa* were associated with disease, whereas those from *Rothia aeria* were not. Conversely, in the lanthipeptide-class-II GCF, *Rothia aeria* BGCs were associated with dental caries, while *Rothia dentocariosa* BGCs were not consistently associated (**Fig. 3**). These patterns highlight that the biosynthetic pathways of specific GCFs, rather than species identity alone, may be critical determinants of oral microbiome function and host disease state. On the other hand, a GCF containing type III PK BGCs displayed mixed health and caries associations: homologous BGCs from *Streptococci* were caries-associated, whereas those from *Leptotrichia* were health-associated (**Fig 3**). This suggests that homologous BGCs from different species can diverge in metabolite structure and function.

**Figure 3:**
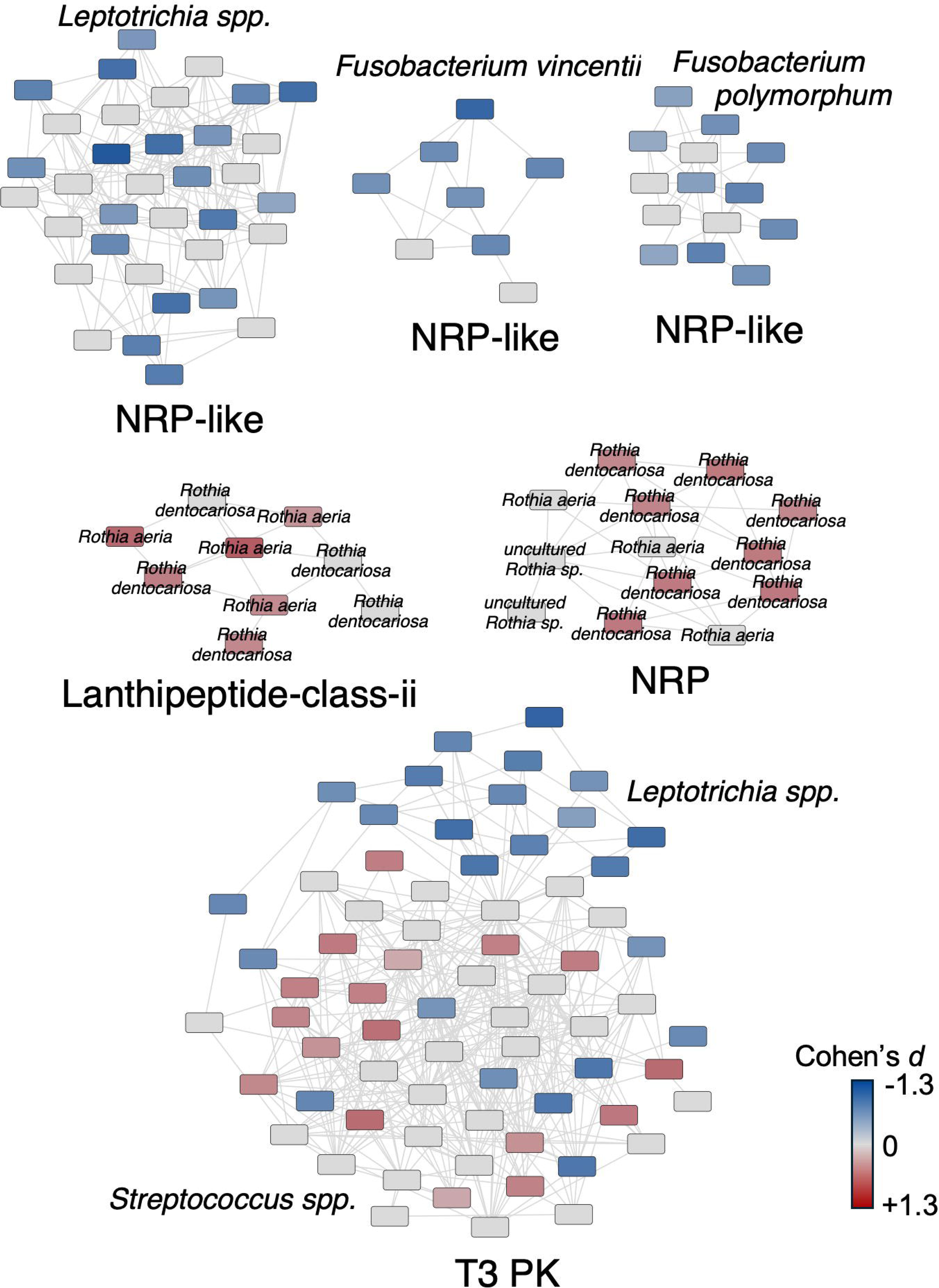
Selected BiG-SCAPE gene cluster families (GCFs) from Dataset 2. Selected gene cluster families from Dataset 2 are shown; see Fig S2 for complete GCF map for Dataset 2. GCFs were grouped using BiG-SCAPE (v2.0) with Pfam-A as the domain database, automatic alignment mode, inclusion of singletons, and a GCF similarity cutoff of 0.5. The resulting similarity network was visualized in Cytoscape using an organic layout, with edges displayed only for distances between 0.3 and 1.0 and filtered to include edge similarity scores (DSS) between 0.5 and 1.0. Nodes represent single biosynthetic gene clusters (BGCs), and edges represent pairwise similarities above the specific thresholds. The GCFs are labeled as their dominant BGC class, and the color gradient indicates the magnitude of each BGC’s Cohen’s *d* value, with blue denoting stronger associations with health and red denoting stronger associations with caries.

### BGC landscape of Dataset 3

Dataset 3 comprised metagenomic samples of the pit and fissure plaque microbiome from 40 adolescents (aged 12 to 13 years), equally divided into 20 with active pit and fissure caries and 20 age-matched, caries-free controls (Pang et al., 2021). Metagenomic libraries were prepared using the NEB Next^®^ Ultra™ DNA Library Prep Kit and sequenced on the Illumina NovaSeq PE150 platform. At the species level, *Actinomyces gerencseriae*, *Prevotella acidifaciens*, *Porphyromonas multisaccharivorax*, *Streptococcus oralis*, *Streptococcus mutans*, and *Parascardovia denticolens* were significantly enriched in the caries-active group. Conversely, *Neisseria elongata*, *Cardiobacterium hominis*, and *Actinomyces johnsonii* were more abundant in caries-free individuals. While *S. mutans* is known as the primary caries-causing pathogen and was significantly higher in the caries-active group, its presence in caries-free subjects and absence in some caries subjects indicated that it alone does not explain all caries cases. Functionally, the caries-active group displayed significant enrichments in carbohydrate metabolism pathways, including the phosphotransferase system, glycolysis/gluconeogenesis, and fructose/mannose metabolism, as well as amino acid metabolism, ABC transporters, and pyrimidine metabolism (Pang et al., 2021).

The metagenomic-assembly analysis on Dataset 3 yielded 8,793 biosynthetic gene clusters (BGCs) that reduced to 2,287 after deduplication. Consistent with the preceding datasets, aryl polyene, RiPP-like/RRE-containing loci, and NRP/NRP-like pathways dominated the major BGC classes (**Fig 1C**). In Dataset 3, terpene BGCs were more likely to be health-associated, whereas disease enriched BGCs mostly included peptide-based pathways (RiPP-like, NRP-PK hybrids, and lanthipeptides). Aryl polyene BGCs were equally distributed between health and disease, similar to Dataset 1 (**Fig 1C**). Uniquely, in Dataset 3, the 20 BGCs with the largest positive Cohen’s *d* values (caries-enriched) contained a more diverse array of natural product classes, including butyrolactone and hybrid NRP–PK encoding BGCs. Specifically, *Streptococcus mutans* contained three RiPP-like BGCs, an NRP-like BGC, and a butyrolactone BGC (**SI Datasheet 1, Tab ‘Dataset 3’**). Aryl polyenes from *Prevotella histicola*, *Prevotella sp*. C561, and *Segatella salivae* were also in the 20 most disease associated BGCs, which showed sequence similarity to the disease-associated aryl polyene GCF identified in Dataset 1. There was also an NRP-PK hybrid BGC from *Propionibacterium acidifaciens*, an NRP-like cluster from *Lautropia dentalis,* and two Type II PK BGCs from *Streptomyces oryzae* and *Actinomycetota*. Among the 20 BGCs with the most negative Cohen’s *d* values (health-enriched), there were several aryl polyenes which belonged to *Cardiobacterium hominis*, several *Neisseria* spp., and *Achromobacter xylosoxidans*. Also included were a lanthipeptide-class-ii BGC from *Corynebacterium matruchotii*, an NRP from *Lautropia dentalis*, an NRP from *Corynebacterium durum,* and a hserlactone from *Bergeriella denitrificans*. From *Kingella*, a RiPP-like cluster from *Kingella oralis* and a terpene from an unidentified *Kingella* sp. were also included. Dataset 3’s diverse BGCs underscore the oral microbiome’s rich biosynthetic potential and the impact of DNA-extraction and sampling methods on observed BGC and species abundance.

BiG-SCAPE networking revealed several GCFs consistently associated with health or disease, several of which were already identified in previous datasets (**Fig. 4, S3**). Notably, the *Prevotella* spp. and *Segatella salivae* aryl polyene GCF that was caries-linked in Dataset 1 was also caries-linked in Dataset 3 (**Fig S3**). Also in Dataset 3 was a GCF with sequence homology to Dataset 1’s largest terpene GCF, but contained BGCs from different species, including *Eikenella corrodens, Neisseria* spp., and *Kingella* spp. (**Fig 4**). This shows that specific BGC functions can be maintained within the oral microbiome, even when carried out by different taxa, suggesting functional redundancy across phylogenetically diverse species. There was also a large RiPP-like GCF with several *Streptococci* species associated with caries and a health-associated NRP-PK hybrid GCF containing BGCs from *Lautropia mirabilis* and *Neisseria cinerea* (**Fig 4**). These examples from Dataset 3 further indicate the consistency and specificity of BGC-associated disease and health correlations. Notably, the reoccurrence of a disease-linked *Prevotella* aryl polyene family and the health-associated *Neisseria* terpene family across independent cohorts from Datasets 1 and 3 shows conserved chemical traits.

**Figure 4:**
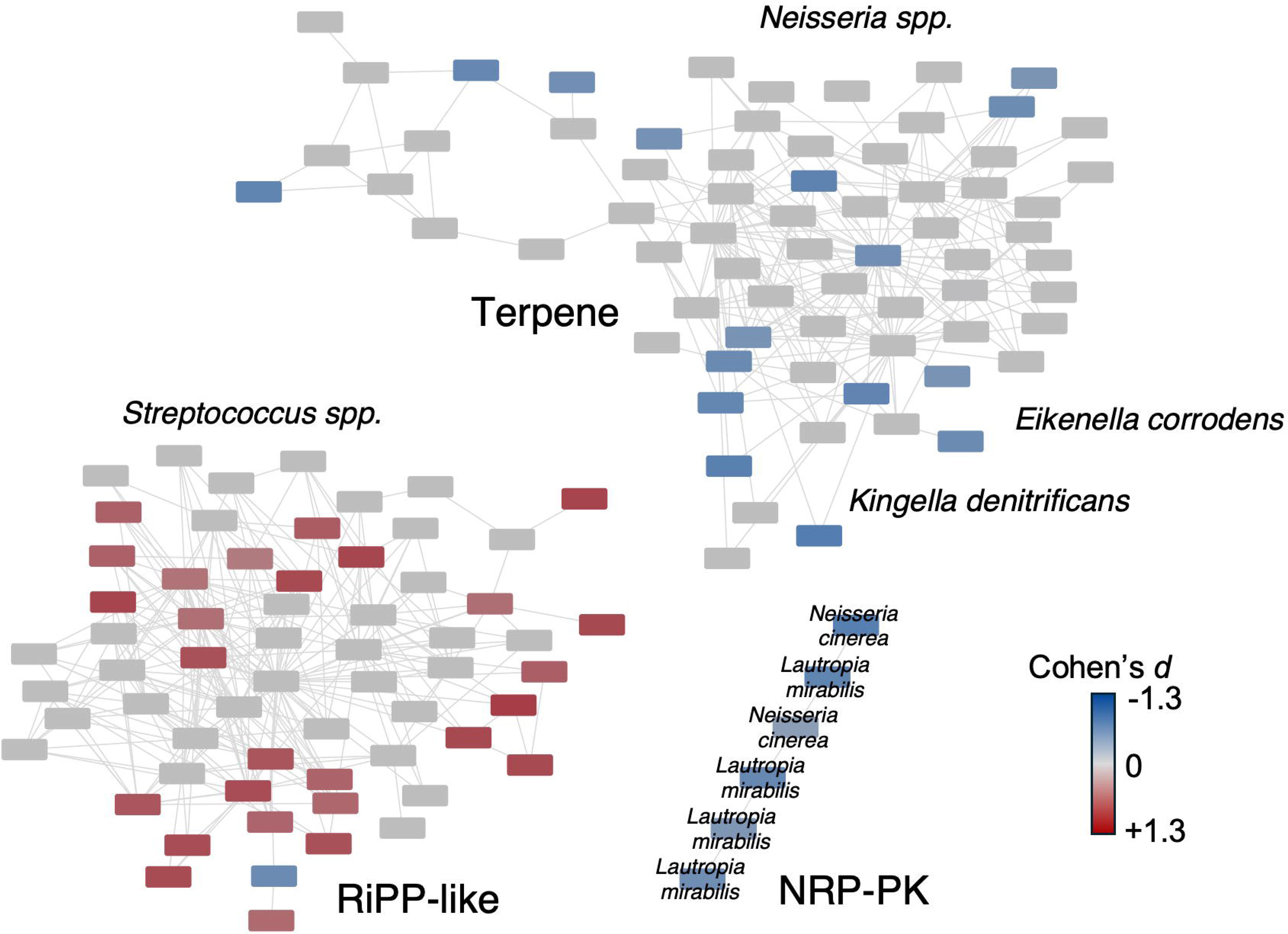
Selected BiG-SCAPE gene cluster families (GCFs) from Dataset 3. Selected gene cluster families from Dataset 3 are shown; see Fig S3 for complete GCF map for Dataset 3. GCFs were grouped using BiG-SCAPE (v2.0) with Pfam-A as the domain database, automatic alignment mode, inclusion of singletons, and a GCF similarity cutoff of 0.5. The resulting similarity network was visualized in Cytoscape using an organic layout, with edges displayed only for distances between 0.3 and 1.0 and filtered to include edge similarity scores (DSS) between 0.5 and 1.0. Nodes represent single biosynthetic gene clusters (BGCs), and edges represent pairwise similarities above the specific thresholds. The GCFs are labeled as their dominant BGC class, and the color gradient indicates the magnitude of each BGC’s Cohen’s *d* value, with blue denoting stronger associations with health and red denoting stronger associations with caries.

### BGC expression landscape from metatranscriptomic dataset

While metagenomics maps genetic potential; metatranscriptomics can capture gene activity, which is important to clarify which genes drive disease and how their expression evolves over time. This analysis is particularly valuable because many BGCs are “cryptic” or known to be activated only under specific conditions. To assess transcriptional activity of the BGCs extracted from Datasets 1–3, we quantified their abundances in a recently published metatranscriptomic dataset comprising supragingival plaque samples from 27 active caries sites and 33 caries-free controls collected from male and female adults aged 18–65 (Mann et al., 2024). All BGCs from Datasets 1-3 were then mapped with BiG-SCAPE based on gene cluster homology, with their associations with health or disease in the metatranscriptomic dataset colored based on their Cohen’s *d* magnitudes.

Based on the alignment of 5,704 BGCs derived from the metagenomic samples in the metatranscriptomic samples, we found that a greater number of aryl polyenes and terpene BGCs were significantly associated with health than dental caries (65% and 90% of significant BGCs, respectively), and NRP and NRP-PK BGCs were more associated with dental caries (90% and 96%, respectively) (**Fig. 5**). Of the 24 BGC classes categorized in this study, 19 contained more BGCs significantly associated with disease than health, suggesting a significant role of microbial specialized metabolism in caries development (**Fig. 5**).

**Figure 5:**
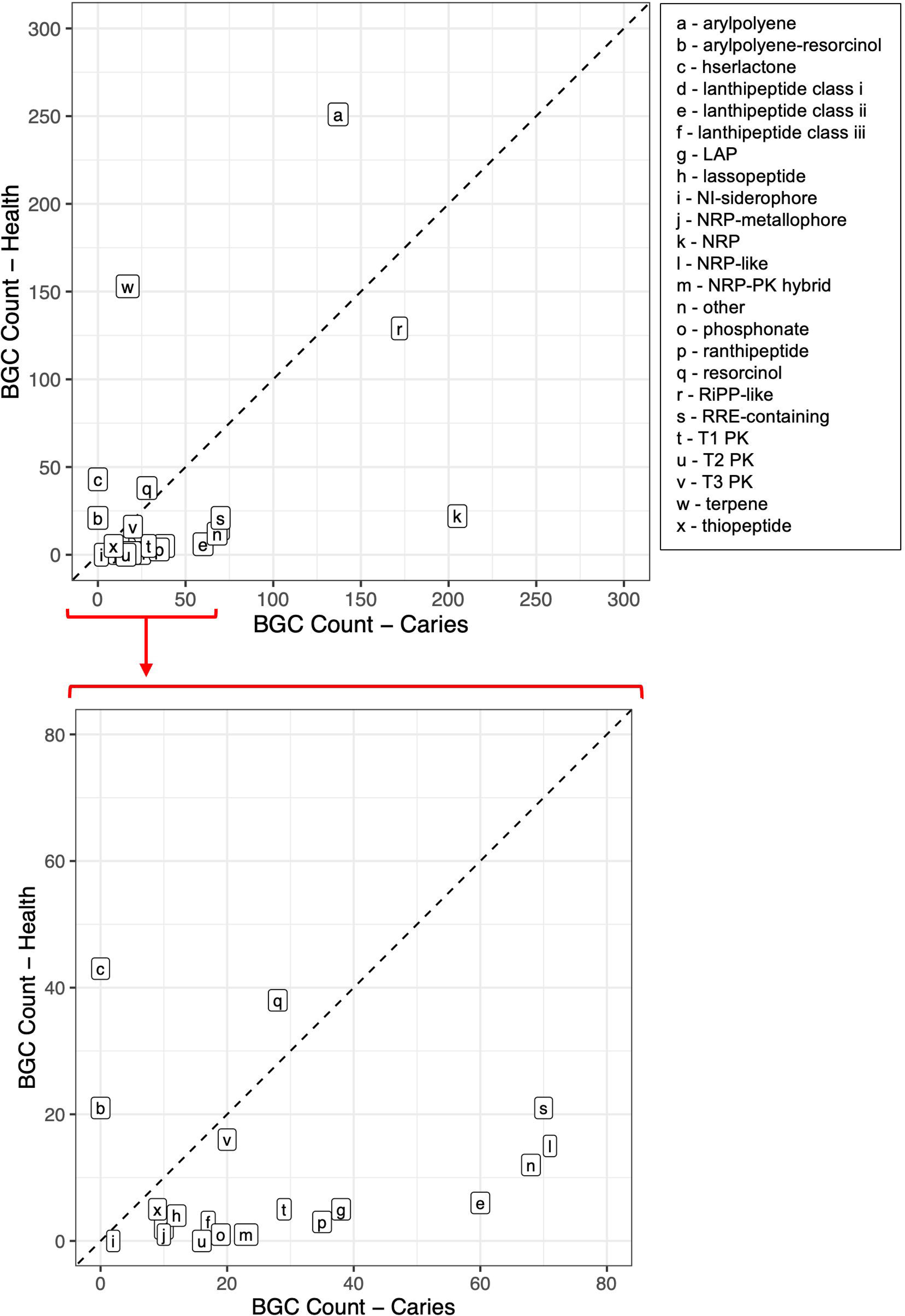
Distribution of biosynthetic gene cluster (BGC) classes in metatranscriptomic dataset. Each square box represents one BGC class, plotted by the number of BGCs significantly enriched in caries samples (x-axis) versus health samples (y-axis). The diagonal dashed line indicates equal representation in both conditions. Letters represent BGC classes and are listed in the right panel. The bottom graph is a zoomed-in version of the top graph from Oto 80 BGC counts.

Our further zoom-in analysis identified that the two major health-associated aryl polyene GCFs extracted in Datasets 1-3 were both associated with health in the metatranscriptomic dataset, which included the *Neisseria* spp. and *Haemophilus parainfluenzae* aryl polyene/resorcinol GCF from Datasets 1 and 3 and the *Capnocytophaga* spp. aryl polyene GCF from Dataset 2 (**Fig S4**, **SI Datasheet 1, Tab ‘Metatranscriptomic’**). This confirmed the prevalence of aryl polyene BGCs in oral health-associated bacteria and their subsequent enrichment in healthy samples. Additionally, the aryl polyene GCF associated with disease in Dataset 3 that contained BGCs from multiple *Prevotella* spp. and *Segatella salivae* was also overexpressed in dental caries samples, indicating its likely role in disease (**Fig. S4**). All terpene BGCs from Datasets 1-3 collapsed into a single GCF and were consistently health-associated in the metatranscriptomic dataset, suggesting that terpenoid biosynthesis may be an indicator of oral health (**Fig. S4**)

In contrast to the large aryl polyene and terpene GCFs, NRP and NRP-PK GCFs were smaller and typically homogenous in a specific species, including those from *Lautropia mirabilis*, *Arachnia propionica*, *Rothia* spp., and *Propionibacterium acidifaciens* (**Fig. S4**). Further, out of the 578 NRP/NRP-PK hybrid singleton BGCs, 191 were association with disease and 19 were associated with health (**SI Datasheet 1, Tab ‘Metatranscriptomic’’**). Complex and modular NRP and NRP-PK BGCs likely make highly potent specialized metabolites to secure niche dominance and drive disease, as demonstrated by a few characterized NRP-PK hybrids from *S. mutans* (Z.-R. Li et al., 2021; Loop Yao et al., 2025).

The number of significant RiPP-like BGCs was similar between health and caries samples in the metatranscriptomic dataset. The major RiPP-like GCFs included: (i) a health-associated RiPP-like/thiopeptide GCF from *Aggregatibacter kilianii*, *Haemophilus influenzae*, and *Mannheimia haemolytica*; (ii) a *Streptococcus* spp. RiPP-like GCF linked to caries; and (iii) a health-associated RiPP-like GCF from *Neisseria lactamica* (**Fig. S5, S6**). The RiPP-like/thiopeptide GCF (i) comprised 50 BGCs, 11 of which were initially health-associated, all of which were from Dataset 2. In the metatranscriptomic dataset, 34 of these BGCs showed a stronger health association than their original metagenomic analysis, with absolute Cohen’s *d* values increasing from <0.5 to 0.5–1.1, and these BGCs originated from all datasets (**Fig. S5, S6**). The *Streptococcus* spp. RiPP-like GCF (ii) drew BGCs from all three datasets and remained caries-associated, though with a weaker effect size than in the metagenomic datasets (Cohen’s *d* < 0.8). Lastly, the *Neisseria lactamica* RiPP-like GCF (iii) contained 61 BGCs, 12 of which were originally associated with health and 30 associated with health and 2 with disease in the metatranscriptomic dataset, with modest absolute Cohen’s *d* values in both the metagenomic and metatranscriptomic datasets (0.3-1). RiPP-like molecules are known for their antimicrobial activities and are likely employed by both health and disease-associated bacteria to control the prevalence of specific bacteria within their niche. These patterns underscore the broad impact of RiPP metabolites across oral microbes in maintaining health and contributing to disease, while structural diversity among them likely reflects species-specific functions.

To identify biosynthetic gene clusters (BGCs) predictive of oral health and disease in the metatranscriptomic dataset, we normalized sample read counts by sequencing depth and applied a centered log-ratio (CLR) transform (Gloor et al., 2017) to account for compositional effects across samples. An elastic-net logistic regression model (α = 0.5) with 10-fold cross-validation achieved an AUROC of 0.798 at λ.1se, indicating strong discriminatory power (Friedman et al., 2010). The λ.1se model retained 45 predictive BGCs (**SI Datasheet 2**), and classification performance was high, with accuracy of 0.966, sensitivity of 0.926, and specificity of 1.000, misclassifying only 2 of 59 samples (**Fig. 6**). Predicted probabilities from the λ.1se model, visualized as jittered points by health versus caries status, showed clear separation around the 0.5 decision threshold (**Fig 6**), consistent with both the confusion matrix and AUROC. Together, these results demonstrate that BGCs can serve as effective predictors of oral health status and highlights their potential as ecological markers of disease.

**Figure 6:**
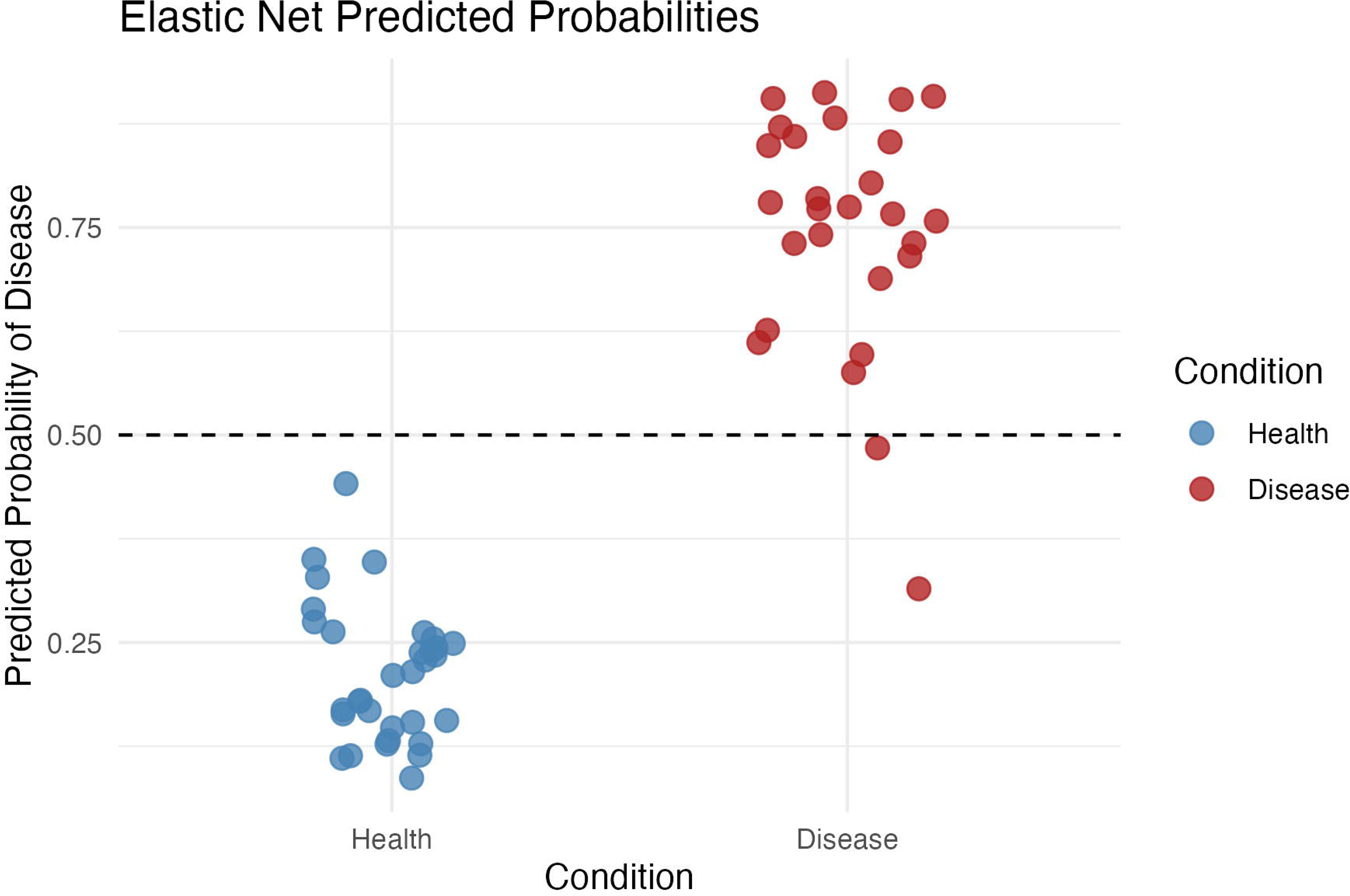
Predicted probabilities from the elastic net model for health- and disease­ associated samples. Predicted probabilities of being classified as health or disease are shown for each metatranscriptomic sample, with points colored by the sample’s true condition (blue= health, red = disease). Probabilities were generated using the l.1 se elastic-net logistic regression model (*a* = 0.5) trained on <u>CLR-transformed</u> BGC expression profiles, which retained 45 predictive BGCs. The dashed horizontal line indicates the 0.5 decision threshold, above which samples are predicted as disease. Health samples cluster below the threshold, while disease samples cluster above it, with only two samples misclassified.

## Discussion

The present study integrated metagenomic assembly, assembly-based BGC prediction, and BGC similarity mapping to investigate how specialized metabolite profiles differ between health and caries-affected oral communities. We identified candidate BGCs and mapped their abundances across three independent metagenomic datasets and one metatranscriptomic dataset, which allowed us to assess their prevalence under health and dental caries conditions. Across datasets, aryl polyenes and terpene pathways were consistently more likely to be enriched and transcriptionally active in health associated samples, whereas NRP and NRP-PK hybrid clusters were disproportionately represented and upregulated in caries.

Aryl polyenes are membrane-associated, carotenoid-like polyenes that represent a widespread but largely uncharacterized class of BGCs across microbial metagenomic surveys (Schöner et al., 2016). Notable examples include flexirubin-type aryl polyenes, originally discovered in the Bacteroides phylum, which can scavenge reactive oxygen species (ROS) and provide photoprotection by delocalizing unpaired electrons along the polyene chain (Schöner et al., 2014). More recently, the aryl polyene biosynthetic pathway of *E. coli* was elucidated and shown to promote oxidative stress tolerance and biofilm formation (Johnston et al., 2021). In the oral cavity, a pilot shotgun metagenomic study comparing health individuals and periodontitis patients found that aryl polyenes were among the most prevalent BGCs in oral samples, consistent with the present study (Koohi-Moghadam et al., 2024).

Terpenoids are among the most characterized classes of natural products, encompassing diverse structures and bioactivities such as environmental stress protection, cell membrane structure and integrity, antimicrobial activity, and host defense (Avalos et al., 2022; González-Burgos & Gómez-Serranillos, 2012). Similar to aryl polyenes, some classes of terpenoids can scavenge free radicals and increase resistance to oxidative bursts (Gutiérrez-del-Río et al., 2021), which may provide protection against salivary peroxidases and host neutrophil-derived ROS in the oral cavity (Zhao et al., 2024). Terpenoids also include hopanoids, which can act as sterol surrogates that rigidify membranes, reduce permeability, and increase tolerance to chemical stresses (Hoshino, 2024). Thus, terpenoids in oral microbes may serve multiple functions, including coping with pH fluctuations, oxidative stress, host immune defenses, and biofilm structure. However, despite being a well-studied natural product class, terpenoids remain predominantly unexplored in the oral cavity and await characterization.

Among disease-associated BGCs, NRP and NRP-PK hybrids encode chemically diverse metabolites that promote interbacterial competition, biofilm development, and stress tolerance (Addington et al., 2024; Fischbach & Walsh, 2006; Wilson et al., 2019). In *Streptococcus mutans,* several NRP-PK hybrids have been identified, including mutanobactin, mutanofactin, mutanoclumpin, and mutanocyclin (Joyner et al., 2010; Z.-R. Li et al., 2021; Loop Yao et al., 2025; Tao et al., 2022). These lipopeptides contribute to biofilm formation, antioxidant defense, and antifungal activity (Barber & Zhang, 2021). Such complex modular peptide pathways confer highly specific functions that drive strain-level dominance and may play analogous roles in other oral pathogens.

Overall, the BGC taxonomy predictions in the present study revealed both “shared” clusters that contain multiple genera and “private” clusters confined to particular species. Shared BGCs, like the aryl polyene BGC family found in *Neisseria* species, suggest some specialized metabolites might serve community-level ecological functions. Conversely, BGCs found in specific species, such as the NRP-PK hybrid from *P. acidifacians,* point to niche-adaptive chemistries that may drive competitive dominance during cariogenesis. The recurrence of homologous BGCs across diverse taxa consistently associated with health or disease suggests that specialized metabolites can be conserved ecological traits, which makes them molecular targets for preventive or therapeutic interventions aimed at maintaining oral microbial homeostasis. Future efforts should prioritize the structural and functional characterization of these metabolites informed by health outcomes and leverage longitudinal multi-omics data to elucidate these metabolites’ dynamic roles during disease progression.

We demonstrate that BGC presence and expression correlate with disease distinctly from the species that harbor these BGCs, and that this observation remains consistent across metagenomic datasets despite variance in sample collection and sequencing strategies. We also show through an elastic-net regression model that BGCs could serve as biomarkers for disease, which could reveal strain level genetic differences without the necessity of strain-level taxonomic analysis. Thus, expanding high-quality metagenomes and metatranscriptomes will be essential to clarify BGC expression patterns across oral niches.

## Methods

### Datasets used in this study

Three metagenomic and one metatranscriptomic datasets were analyzed in this study. Paired-end reads were downloaded from the NCBI Sequence Read Archive (SRA) under BioProjects PRJNA766357, PRJNA383868, PRJNA380711, and PRJEB60355. Specific accession numbers of all samples used in this study are included in the Supplementary Information.

### Sequence quality control and metagenomic assembly

Adapter sequences were removed with Trimmomatic v0.39 (Bolger et al., 2014), allowing up to two mismatches in the seed region and applying clip-score thresholds of 30 (palindrome) and 10 (simple) while retaining orphaned mates for performance computing cluster operated by Berkeley Research Computing at the University of California, Berkeley.

### Biosynthetic gene cluster prediction and annotation

Metagenomic assemblies were screened for biosynthetic gene clusters with antiSMASH v7.0 (Blin et al., 2023). The following annotation modules were enabled: KnownClusterBlast, ClusterBlast subclusters, transcription-factor binding sites (TFBS), Active-Site Finder (ASF), and RiPP Recognition Element (RRE). Open reading frames were called with the Prodigal gene-finding algorithm to ensure consistent coding-sequence boundaries. Resulting GenBank-formatted output files for every sample were consolidated into a single directory to facilitate downstream comparative analyses of BGCs.

### Construction of non-redundant BGC database

All antiSMASH-predicted BGCs were compared pairwise with MultiGeneBlast v1.1.14 (Medema et al., 2013) to identify and collapse redundant biosynthetic gene clusters (BGCs). For each dataset, every BGC was queried against a database containing all the other BGCs from that same dataset, yielding an all-vs-all comparison. Similarity searches required (i) ≥ 80 % sequence coverage of the query region and (ii) ≥ 90 % nucleotide identity across aligned segments; searches were restricted to the annotated BGC coordinates to avoid spurious hits in flanking DNA.

MultiGeneBlast HTML and tabular outputs were exported for downstream parsing. Redundancy was assessed with a custom Python script (repository link in GitHub). For any BGC pair, a cluster was flagged as a duplicate when at least 90 % of its genes were matched by BLAST hits that each exceeded 90% identity, and the partner BGC contained more genes than were matched (i.e., the putative duplicate was a strict subset of a larger cluster). Duplicates were removed, leaving a single representative for every unique BGC. The resulting non-redundant BGC list served as the input for all subsequent occurrence analyses.

### Mapping reads to biosynthetic gene clusters

All non-redundant BGC sequences were exported from their GenBank files as nucleotide FASTA records and concatenated into a single reference file. This reference was indexed with BWA-MEM2 v2.2.1 (Vasimuddin et al., 2019), after which paired-end reads from each unassembled metagenomic and metatranscriptomic sample were aligned to the index. Mapping jobs used the MEM algorithm with the number of CPU threads set to the node allocation (24) and a minimum alignment score threshold of 80, ensuring that only confidently matched reads were retained. Alignments were written in SAM format and parsed to tally reads mapping to each BGC in every sample, generating a table of per-sample BGC counts. BGC counts were then normalized based on each sample’s reads.

### Gene-cluster family classification and network visualization

To group the non-redundant BGC list into gene-cluster families (GCFs), sequences were analysed with BiG-SCAPE (v2.0.0b6) (Navarro-Muñoz et al., 2020). Domain annotation relied on the Pfam-A HMM library implementing and the auto alignment mode; singletons were retained so that unique clusters were not lost. GCFs were defined at a similarity cut-off of 0.5, and the mix distance metric (which averages the “anchor”, “domain”, and “sequence similarity” scores) balanced sensitivity and specificity across diverse BGC classes.

The resulting network file was imported into Cytoscape v3.10 (Shannon et al., 2003) for visual exploration. Edges were filtered to retain only those with a BiG-SCAPE distance between 0.3 and 1.0 and a domain sequence similarity (DSS) score between 0.5 and 1.0, producing a manageable graph that highlights the most relevant inter-cluster relationships while excluding weak or spurious links.

### Species-level annotation of biosynthetic gene clusters

To infer the likely taxonomic origin of each BGC, we first extracted the largest core biosynthetic protein (ketosynthase, adenylation, condensation, or equivalent “anchor” enzyme) from every cluster using a custom Python script (extract_anchors_2.py). These anchor sequences were searched against the NCBI ClusteredNR database with NCBI BLAST+ v2.16.0 (BLASTp task) (Camacho et al., 2009). Searches used an E-value threshold of 1 × 10 and reported up to 100 subject sequences per query while limiting each subject to a single highest-scoring segment pair. The output table was then sorted by query identifier, descending bitscore, and ascending E-value; the top-ranking hit for each anchor protein was retained to assign a putative species designation to its corresponding BGC.

### Statistical analysis and effect-size estimation

All statistics were carried out in RStudio 2023.03.1 + 446 running R 4.3.2. Data wrangling and visualization employed the tidyverse meta-package v2.0.0 (Wickham et al., 2019). For pairwise comparisons between health and dental-caries groups, BGC read-counts were assessed with a two-sided Wilcoxon rank-sum test (exact calculation disabled); significance was defined as *p* < 0.05. Practical effect sizes were quantified as Cohen’s *d*, calculated as the difference between the group means (caries – healthy) divided by the pooled standard deviation. Mean values, *p*-values, and effect sizes were exported to tab-separated summary tables for downstream interpretation and figure generation.

## Supporting information

SI Datasheet 1

SI Datasheet 3

SI Datasheet 2

## Acknowledgements

We thank Shiyi Yang (Computational biology, UC Berkeley) for assistance with biostatistical analysis and Aarav Shah, Arun Kamath, and Steven Yu (College of Computing, Data Science, and Society, University of California, Berkeley) for their early work on the computational pipeline.

## Data Availability

The authors declare that all the data supporting the findings of the present study are available within the manuscript, the Supplementary Information, and/or in the following public data repositories. The metagenomic and metatranscriptomic datasets can be accessed in NCBI’s database, with BioProjects PRJNA766357, PRJNA383868, PRJNA380711, and PRJEB60355. All sample details used in this study can be found in SI Datasheet 3.

