## Supplementary Figures for "The biosynthetic gene cluster landscape of the oral microbiome across health and dental caries"

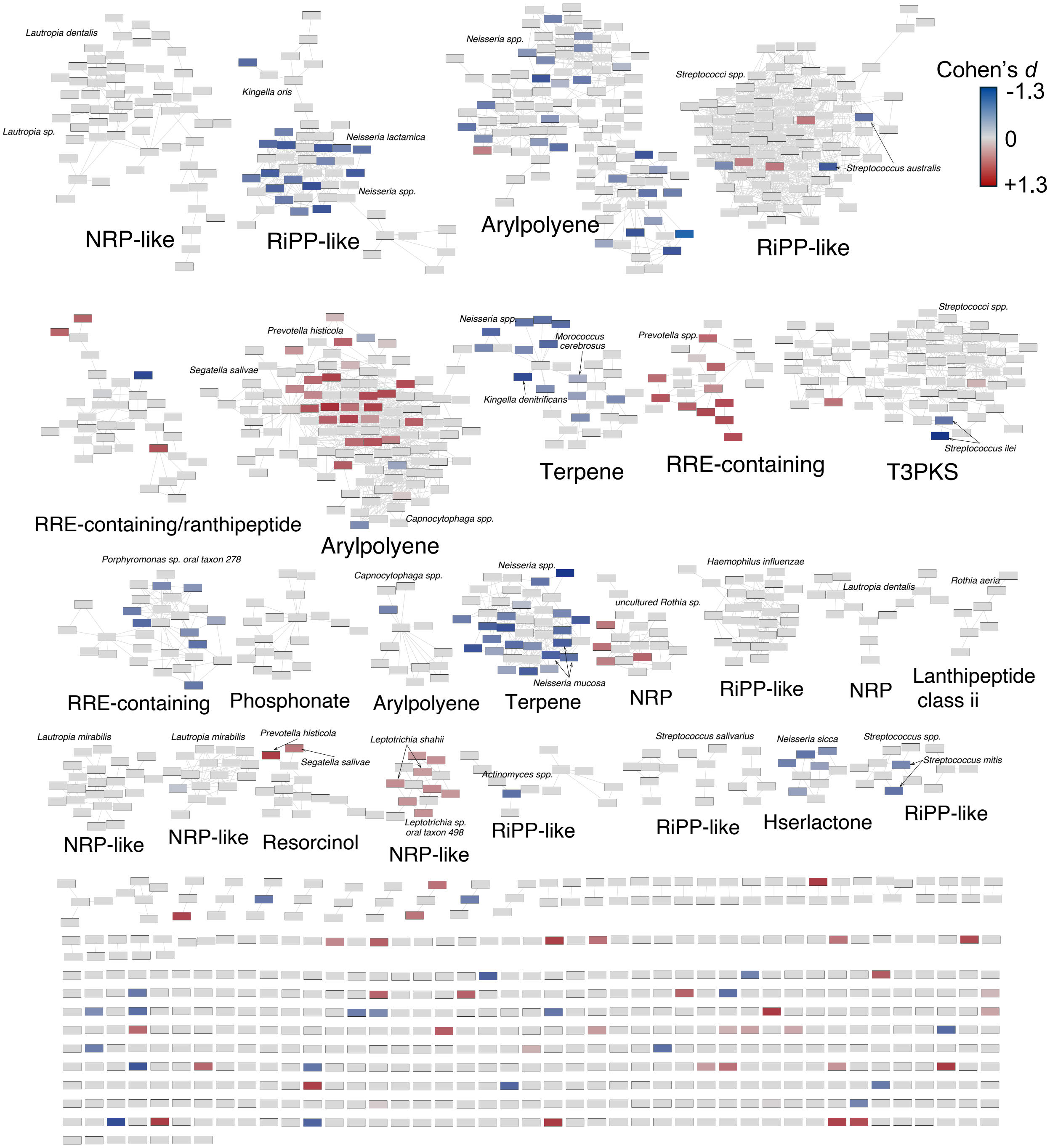


**Figure S1: BiG-SCAPE gene cluster family (GCFs) network analysis from Dataset 1.** Gene clusters from Dataset 1 were grouped into gene cluster families (GCFs) using BiG-SCAPE (v2.0) with Pfam-A as the domain database, automatic alignment mode, inclusion of singletons, and a GCF similarity cutoff of 0.5. The resulting similarity network was visualized in Cytoscape using an organic layout, with edges displayed only for distances between 0.3 and 1.0 and filtered to include edge similarity scores (DSS) between 0.5 and 1.0. Nodes represent individual biosynthetic gene clusters, and edges represent pairwise similarities above the specific thresholds. Selected GCFs are labeled by their BGC class. The color gradient indicates the magnitude of each BGC's Cohen's *d* value, with blue denoting stronger associations with health and red denoting stronger associations with disease.

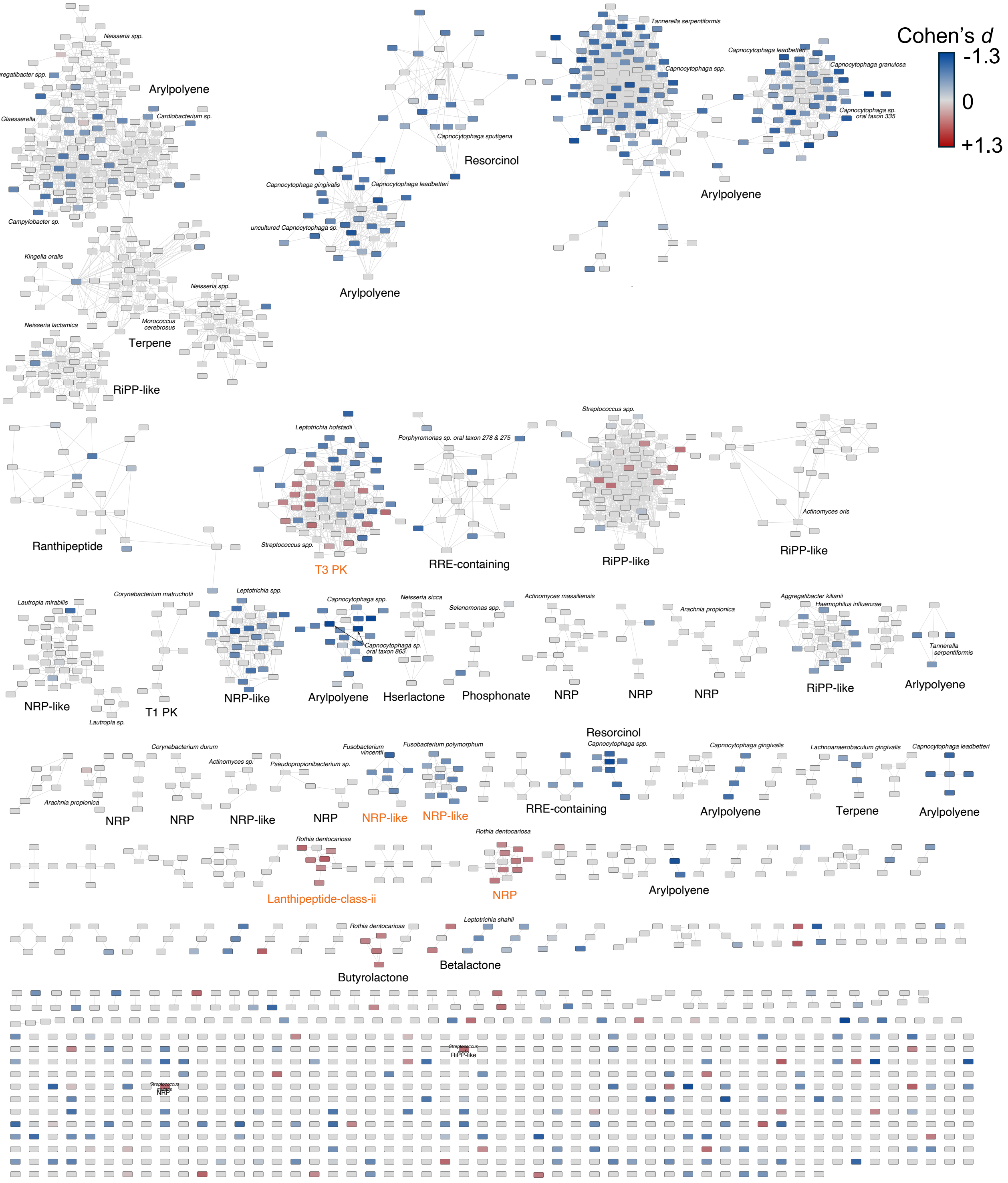


**Figure S2: BiG-SCAPE gene cluster family (GCFs) network analysis from Dataset 2.** Gene clusters from Dataset 2 were grouped into gene cluster families (GCFs) using BiG-SCAPE (v2.0) with Pfam-A as the domain database, automatic alignment mode, inclusion of singletons, and a GCF similarity cutoff of 0.5. The resulting similarity network was visualized in Cytoscape using an organic layout, with edges displayed only for distances between 0.3 and 1.0 and filtered to include edge similarity scores (DSS) between 0.5 and 1.0. Nodes represent individual biosynthetic gene clusters, and edges represent pairwise similarities above the specific thresholds. Selected GCFs are labeled by their BGC class. The color gradient indicates the magnitude of each BGC's Cohen's *d* value, with blue denoting stronger associations with health and red denoting stronger associations with disease. Orange GCF labels correspond to selected GCFs in Figure 3.

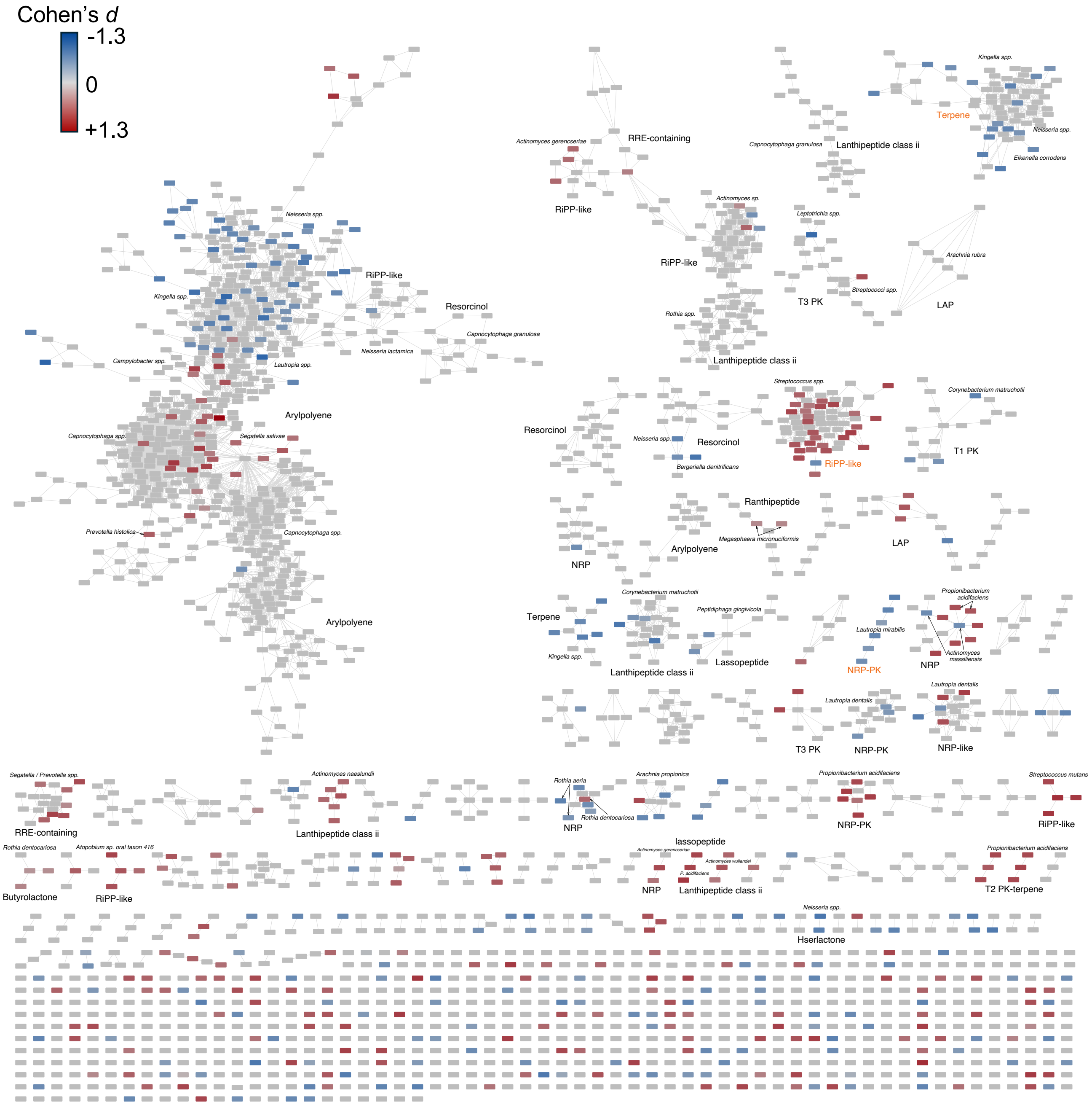


**Figure S3: BiG-SCAPE gene cluster family (GCFs) network analysis from Dataset 3.** Gene clusters from Dataset 3 were grouped into gene cluster families (GCFs) using BiG-SCAPE (v2.0) with Pfam-A as the domain database, automatic alignment mode, inclusion of singletons, and a GCF similarity cutoff of 0.5. The resulting similarity network was visualized in Cytoscape using an organic layout, with edges displayed only for distances between 0.3 and 1.0 and filtered to include edge similarity scores (DSS) between 0.5 and 1.0. Nodes represent individual biosynthetic gene clusters, and edges represent pairwise similarities above the specific thresholds. Selected GCFs are labeled by their BGC class. The color gradient indicates the magnitude of each BGC's Cohen's *d* value, with blue denoting stronger associations with health and red denoting stronger associations with disease. Orange GCF labels correspond to selected GCFs in Figure 4.

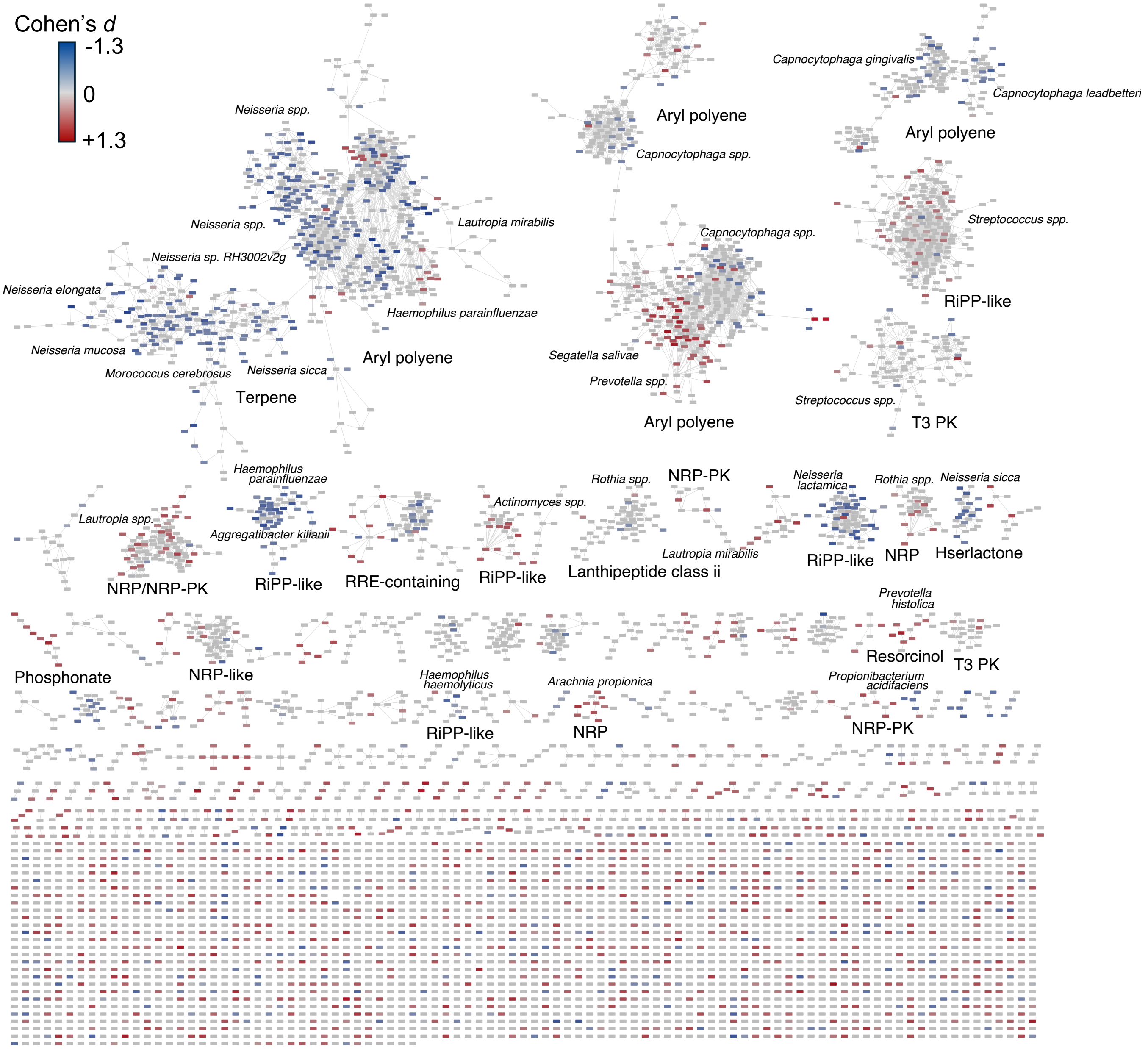


**Figure S4: BiG-SCAPE gene cluster family (GCFs) network analysis from metatranscriptomic dataset.**

Gene clusters from the metatranscriptomic dataset were grouped into gene cluster families (GCFs) using BiG-SCAPE (v2.0) with Pfam-A as the domain database, automatic alignment mode, inclusion of singletons, and a GCF similarity cutoff of 0.5. The resulting similarity network was visualized in Cytoscape using an organic layout, with edges displayed only for distances between 0.3 and 1.0 and filtered to include edge similarity scores (DSS) between 0.5 and 1.0. Nodes represent individual biosynthetic gene clusters, and edges represent pairwise similarities above the specific thresholds. Selected GCFs are labeled by their BGC class. The color gradient indicates the magnitude of each BGC's Cohen's *d* value, with blue denoting stronger associations with health and red denoting stronger associations with disease.

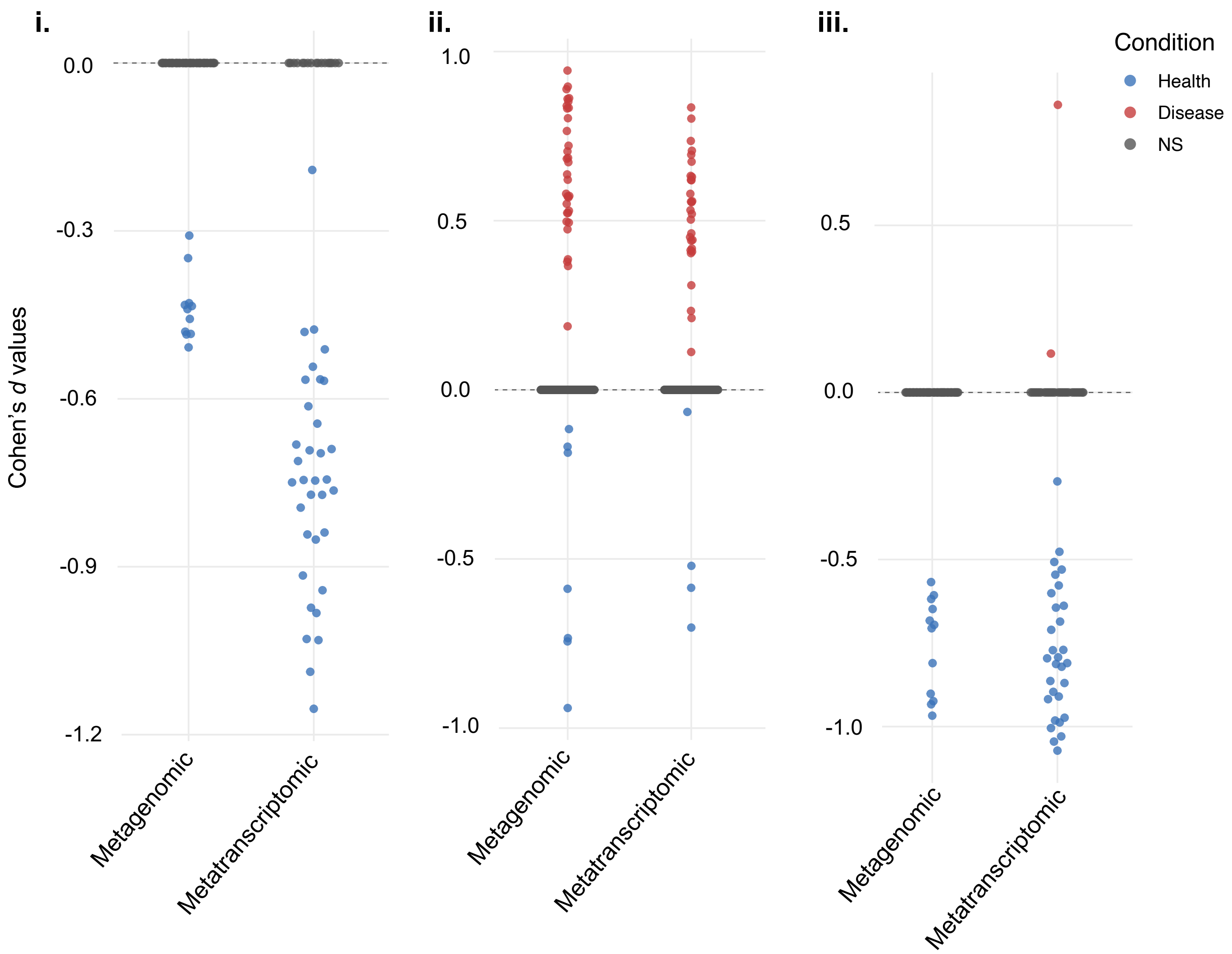


**Figure S5: Jitter plot of RiPP- like gene cluster families across datasets.** Three major RiPP-like gene cluster families (GCFs) (i-iii) are plotted according to their Cohen's *d* values across Datasets 1–3 and the metatranscriptomic dataset. Data points correspond to individual BGCs classified as significantly associated with health (blue), caries (red), or not significant (grey).

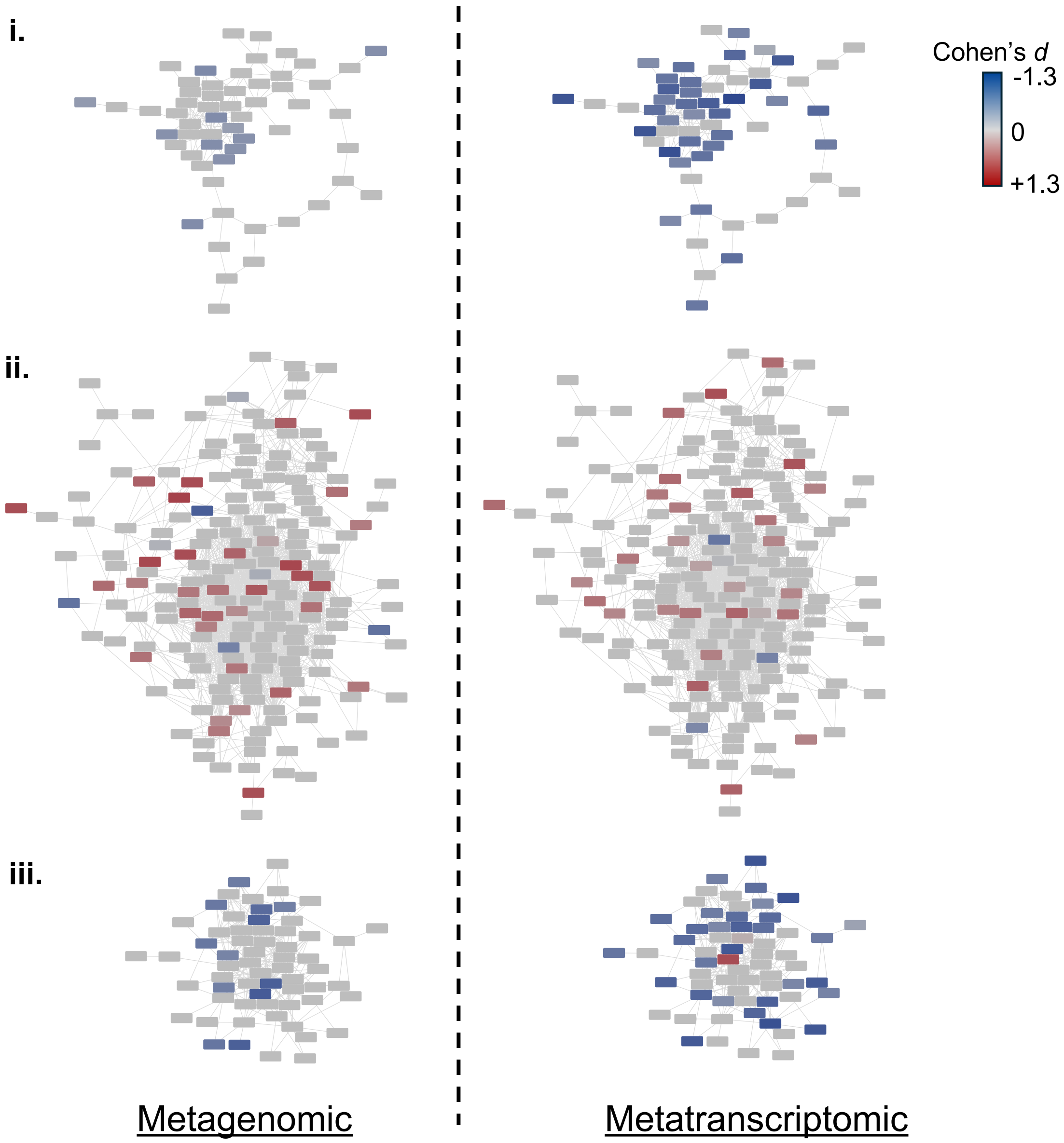


**Figure S6: Gene cluster families of RiPP-like BGCs across datasets.** Three major RiPP-like gene cluster families (GCFs; i–iii) are displayed with nodes representing individual BGCs. In the left panel, BGCs are colored by their Cohen's *d* values from Datasets 1–3; in the right panel, the same BGCs are colored by Cohen's *d* values from the metatranscriptomic dataset. The color gradient spans red (caries-associated) to blue (health-associated), with non-significant BGCs shown in grey.
